# Curcumin extends the lifespan of aging postmitotic cells with mitochondrial dysfunction

**DOI:** 10.1101/2023.09.06.556469

**Authors:** Arshia Naaz, Yizhong Zhang, Nashrul Afiq Faidzinn, Sonia Yogasundaram, Mohammad Alfatah

**Affiliations:** Genome Institute of Singapore (GIS), Agency for Science, Technology and Research (A*STAR), 60 Biopolis Street, Genome #02-01, Singapore 138672, Republic of Singapore; Bioinformatics Institute (BII), Agency for Science, Technology and Research (A*STAR), 30 Biopolis Street, Matrix #07-01, Singapore 138671, Republic of Singapore

**Keywords:** Curcumin, postmitotic cellular aging, lifespan, healthspan, mitochondria, TORC1, yeast, human

## Abstract

Aging is an inevitable biological process intricately linked to age-related diseases, including cardiovascular diseases, neurodegeneration, sarcopenia, and age-related macular degeneration. These ailments are often exacerbated by mitochondrial dysfunction, which plays a pivotal role in postmitotic cells. Curcumin, a natural compound, is explored for its anti-aging potential. This study explores the influence of curcumin on the postmitotic cellular lifespan (PoMiCL) of yeast during chronological aging, examining its potential implications for age-related diseases. Our findings reveal that curcumin significantly extends the lifespan of postmitotic wildtype yeast cells, with maximal effects observed at lower concentrations, displaying a hormetic response. Importantly, curcumin mitigates accelerated aging in cells afflicted by mitochondrial dysfunction. Intriguingly, the hormetic effect is absent under these conditions. Mechanistically, curcumin enhances ATP levels but induces oxidative stress and inhibits TORC1. These findings shed light on curcumin’s potential as an anti-aging modulator and its relevance to age-related diseases, offering insights into novel therapeutic approaches for healthy aging while highlighting the context-dependent nature of its effects.

## INTRODUCTION

Aging, a natural biological phenomenon that occurs as time passes, is a universal occurrence observed in all living organisms. It involves a gradual deterioration in cellular functions and is closely connected to the emergence of various age-related diseases ^1–3^. These illnesses not only present substantial hurdles for the healthcare system but also affect the well-being of aging individuals. With the world’s population aging at an increasing rate ^4^, comprehending the mechanisms behind aging and investigating approaches to mitigate its consequences have become crucial objectives in biomedical research.

One key factor associated with age-related diseases is mitochondrial dysfunction ^5–8^. Mitochondria, often referred to as the “powerhouses of the cell,” play a central role in energy production, cellular metabolism, and the regulation of oxidative stress ^9^. However, with advancing age, mitochondrial function can deteriorate, leading to a cascade of events that contribute to the pathogenesis ^5–9^. The association between mitochondrial dysfunction in postmitotic cells and the aging process, as well as its implications for cardiovascular diseases (CVDs), neurodegenerative diseases (NDDs), sarcopenia, and age-related macular degeneration (AMD), represents a pivotal area of research in the field of aging biology. Mitochondrial dysfunction has been linked to CVDs, where impaired energy production and increased oxidative stress play critical roles in the development of cardiac pathologies ^10–12^. It is also a hallmark of NDDs, including Alzheimer’s and Parkinson’s diseases, where mitochondrial dysfunction exacerbates neuronal damage and cell death ^13,14^. Moreover, mitochondrial dysfunction has been implicated in sarcopenia (age-related loss of muscle mass and function) ^15,16^, as well as AMD, a leading cause of vision loss in the elderly ^17–19^.

In the quest for effective anti-aging interventions, natural compounds have emerged as promising candidates ^20,21^. Recent research has drawn attention to the potential impact of curcumin, a polyphenolic bioactive compound derived from the rhizome of Curcuma longa (turmeric), for its therapeutic potential across a wide spectrum of diseases including mitigating the effects aging and increasing lifespan ^22–25^. However, a critical knowledge gap exists in the realm of real-world observational data that examines the impact of dietary curcumin, primarily sourced from the consumption of curry in food, on health and longevity.

Two groundbreaking observational cohort studies conducted among middle-aged and older Asian adults living in Singapore have revealed an intriguing connection between the consumption of curcumin-rich foods and healthspan ^26,27^. This study represents the inaugural longitudinal exploration of the cognitive advantages linked to curcumin obtained from natural dietary sources in human subjects. The results emphasize the potential health and longevity-enhancing effects of curcumin in the diet from natural sources, providing valuable insights into how consuming curry may affect various health aspects, including cognitive benefits over time and the potential to extend the lifespan of patients with cardio-metabolic and vascular diseases.

The present study elucidates the molecular and cellular mechanisms by which curcumin influences the postmitotic cellular lifespan (PoMiCL) within the context of chronological aging, particularly when mitochondrial dysfunction is a contributing factor. Understanding how curcumin affects PoMiCL during chronological aging, especially in the presence of mitochondrial dysfunction, is crucial, as it may offer insights into novel strategies for promoting healthy aging and combating age-related diseases, including CVDs, NDDs, sarcopenia, and AMD.

We conducted a series of thoroughly designed PoMiCL experiments in yeast, a well-established model organism for aging research ^28–36^. These assays allowed us to systematically explore curcumin’s influence on cellular survival at various time points during chronological aging. Our mechanistic findings shed light on the complex interplay between curcumin treatment, mitochondrial function, and PoMiCL, offering valuable insights into the potential utility of curcumin as an anti-aging modulator. By uncovering the intricacies of curcumin’s effects on postmitotic cells, we provide valuable insights that may ultimately contribute to the development of novel therapeutic approaches for promoting longevity and enhancing the quality of life in an aging world.

## RESULTS

### Curcumin extends postmitotic cellular lifespan (PoMiCL) during chronological aging

We investigated the effect of curcumin treatment on the lifespan of yeast wildtype non-dividing cells within the context of chronological aging. We performed PoMiCL assays that provided us with a dynamic platform to meticulously dissect the impact of curcumin on cellular survival during the gradual progression of chronological aging. To initiate the experimental regimen, yeast wildtype cells were systematically exposed to a spectrum of curcumin concentrations, ranging from 12.5 µM to 200 µM, within the confines of a 96-well microplate. To initiate the chronological aging process, we first allow the cells to transition into the postmitotic state. This was achieved by cultivating the cell cultures in medium until they reach the stationary phase, which marking the beginning of the PoMiCL assays during chronological aging. Subsequently, we embarked on an exploration of cellular responses at different time points, including day 3, day 6, day 13, and day 20, post-initiation of curcumin treatment. Notably, the viability of postmitotic cells was quantified through a comprehensive assessment of outgrowth, a metric that was precisely normalized in relation to the baseline data obtained on day 3 (Figures 1A – 1D).

**Figure 1.**
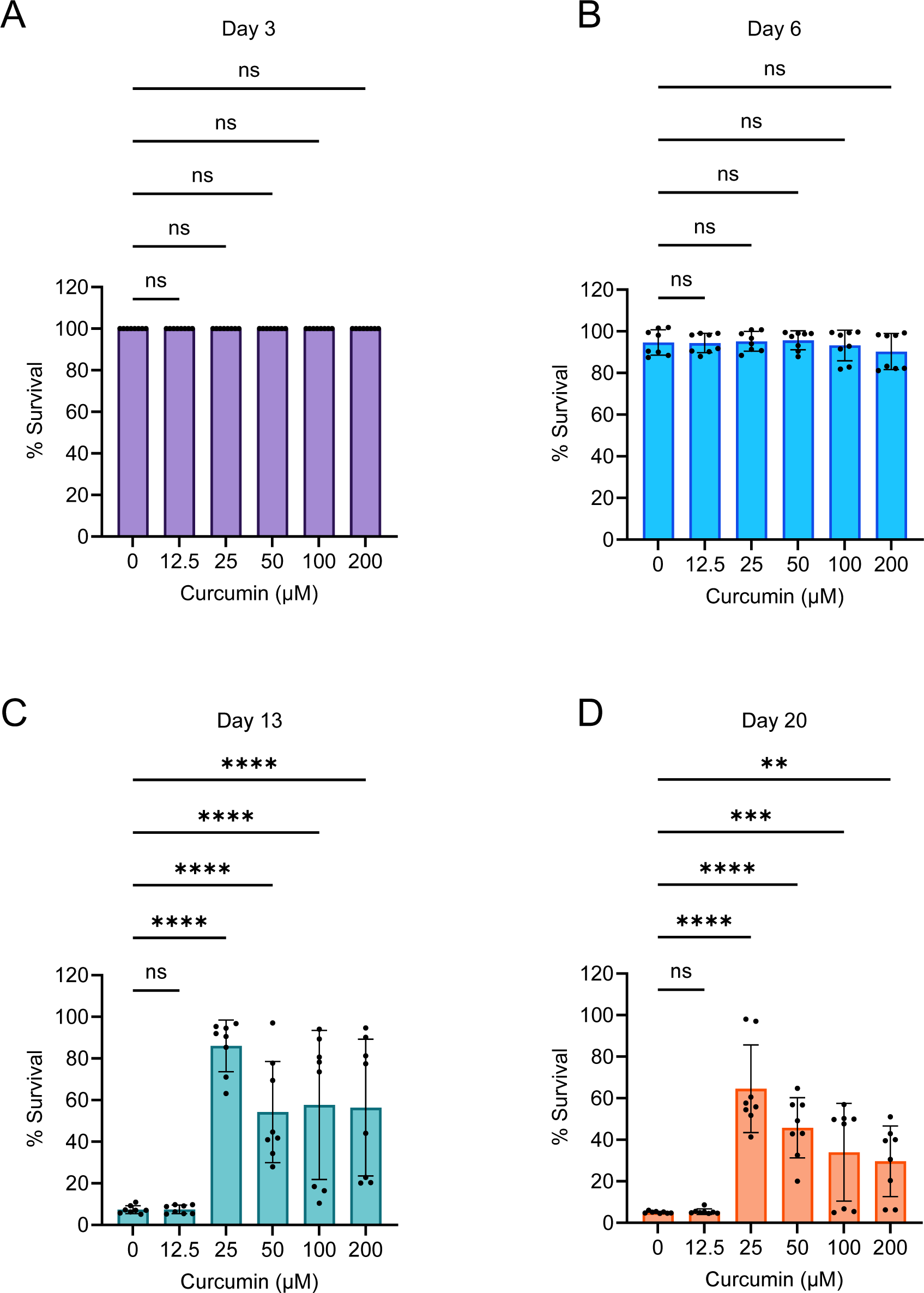
Curcumin increases the postmitotic cellular lifespan (PoMiCL) of yeast during chronological aging. The effect of curcumin treatment on the PoMiCL of wildtype yeast was evaluated in an SD medium using a 96-well plate. Cells were incubated with indicated concentrations of curcumin in the 96-well plate. The survival of chronological aging postmitotic cells was measured on (A) day 3, (B) day 6, (C) day 13 and (D) day 20, relative to the outgrowth of day 3. The data are presented as means ± SD (n=8). Statistical significance was determined as follows: **P < 0.01, ***P < 0.001, ****P < 0.0001 and ns (non-significant), based on a two-way ANOVA followed by Dunnett’s multiple comparisons test.

Upon conducting a thorough PoMiCL analysis, our observations revealed a lack of discernible differences in cell survival on day 6, where both curcumin-treated and untreated postmitotic cells exhibited approximately 100% survival rates (Figure 1B). However, a notable contrast emerged on day 13. Aging postmitotic cells subjected to curcumin treatment, specifically at a concentration of 25 µM, displayed a striking enhancement in their survival rates, increasing to approximately 90%. In stark contrast, cells within the control group, devoid of curcumin treatment (receiving only DMSO), demonstrated a miserable survival rate, falling below the 10% threshold (Figure 1C). The divergence in survival dynamics persisted, substantiated by our observations on day 20, wherein curcumin-treated postmitotic cells maintained a robust survival rate of approximately 65% (Figure 1D).

These empirical findings robustly endorse the hypothesis that curcumin possesses the remarkable capacity to forestall cellular aging and, consequently, extend the PoMiCL of yeast. Our outcomes are notably consistent with an amount of prior research that has underscored the beneficial effects of curcumin in the context of cellular longevity ^37–39^. Furthermore, our novel insights into curcumin’s influence on cellular lifespan complement with analogous investigations conducted using different genetic backgrounds and distinct cell survival assays ^40^. Collectively, these results compellingly advocate for the potential utility of curcumin as a modulator of cellular aging and lifespan extension across diverse eukaryotic species.

Intriguingly, our investigations unveiled a nuanced dosage-dependent phenomenon. The maximal benefits of curcumin with regard to cellular lifespan extension were achieved at lower concentrations, whereas higher concentrations demonstrated diminishing efficacy (Figures 1C and 1D). This intriguing observation suggests to a biphasic dose-response relationship, indicative of a hormetic effect exerted by curcumin on cellular lifespan. To confirm our findings, we conducted the PoMiCL assay using human kidney cells (HEK293) ^41^. We observed a similar trend to that in yeast, wherein lower concentrations of curcumin exhibited a more effective anti-aging activity compared to higher concentrations (Figure S1). This result indicates that the anti-aging properties of curcumin are functionally consistent across yeast and human cells, showing an association with hormetic effects.

It’s worth noting that hormetic effects of curcumin have been previously documented in various biological contexts ^42^. For instance, previous studies have explored the wound-healing capabilities of human skin fibroblasts during the aging process ^43^. Thus, our findings not only contribute to the growing body of evidence supporting the beneficial effects of curcumin but also advance our understanding of hormesis in the context of cellular longevity during the process of chronological aging.

### Curcumin extends PoMiCL with mitochondrial function deficiencies

In our previous study, we demonstrated that inducing mitochondrial dysfunction through chemical or genetic means accelerates cellular aging and reduces the lifespan of postmitotic cells ^44^. Given the pivotal role of mitochondrial function in age-related diseases and the potential influence of curcumin on cellular aging, we explored the impact of curcumin on the lifespan of postmitotic cells deficient in mitochondrial function during chronological aging. We assessed the effects of curcumin on the PoMiCL of mitochondrial gene deletion mutants associated with the loss of respiratory activity and energy synthesis. *PET100* (*YDR079W*) is a yeast gene critical for assembling and maintaining the mitochondrial respiratory chain complex IV (cytochrome c oxidase), which is essential for mitochondrial ATP production and overall cellular energy metabolism ^45^. We conducted PoMiCL experiments on *pet100Δ* mutants treated with various curcumin concentrations and compared them with wildtype cells. The survival of aging postmitotic cells was measured at specified time intervals relative to the outgrowth on day 3 (Figures 2A – 2D).

**Figure 2.**
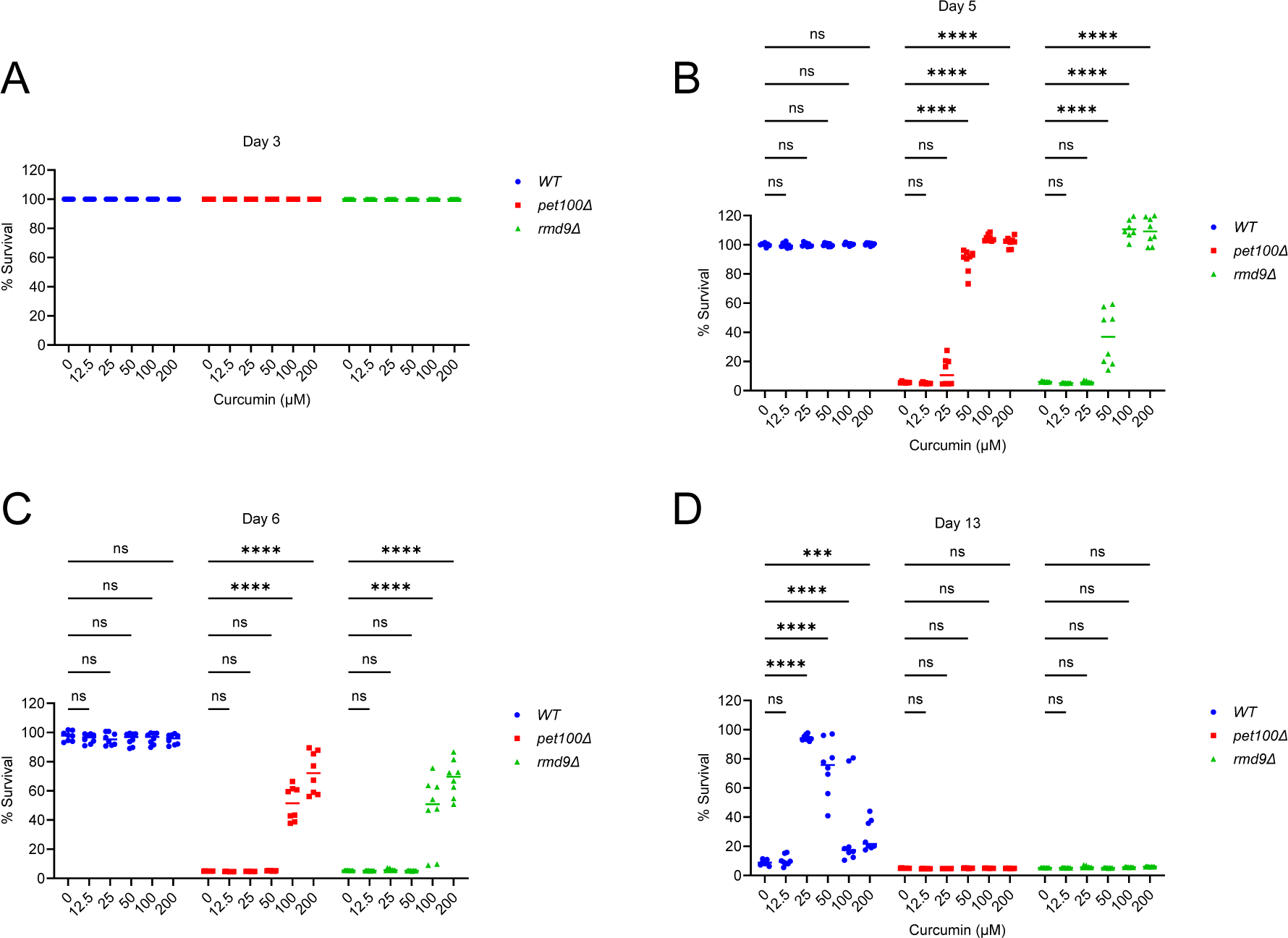
Curcumin increases the postmitotic cellular lifespan (PoMiCL) of yeast with mitochondrial function deficiencies. The effect of curcumin treatment on the PoMiCL of yeast wildtype and mitochondrial dysfunction mutants (*pet100Δ* and *rmd9Δ*) was evaluated in an SD medium using a 96-well plate. Cells were incubated with indicated concentrations of curcumin in the 96-well plate. The survival of chronological aging postmitotic cells was measured on (A) day 3, (B) day 5, (C) day 6 and (D) day 13, relative to the outgrowth of day 3. The data are presented for replicates (n=8). Statistical significance was determined as follows: ***P < 0.001, ****P < 0.0001 and ns (non-significant), based on a two-way ANOVA followed by Dunnett’s multiple comparisons test.

On day 5 and day 6, the survival of *pet100Δ* mutant (DMSO control) was below 10% (Figures 2B and 2C). Conversely, at the same chronological aging time points (day 5 and day 6), the survival rate of wildtype (DMSO control) remained at approximately 100% (Figures 2B and 2C). These results align with our previous findings, illustrating that mitochondrial dysfunction leads to accelerated aging and a shortened PoMiCL ^44^. Remarkably, curcumin treatment prevents accelerated aging and enhances the PoMiCL of mitochondrial dysfunction *pet100Δ* mutant (Figures 2B and 2C). The survival rate of *pet100Δ* cells treated with curcumin (50 µM) was around 90%, in contrast to less than 10% in untreated cells (Figure 2B). Interestingly, the survival rate increased with higher concentrations (100 µM and 200 µM) of curcumin-treated cells, achieving nearly 100% viability on day 5 (Figure 2B). A similar concentration-dependent trend was observed in increased lifespan on day 6. The survival rate of *pet100Δ* cells treated with the minimal effective curcumin concentration (100 µM) was approximately 60%, compared to less than 10% in untreated cells (Figure 2C). A higher curcumin concentration (200 µM) further boosted the survival of *pet100Δ* cells to roughly 80% on day 6 (Figure 2C). These findings indicate that curcumin represents a promising anti-aging intervention that mitigates accelerated cellular aging and prolongs the lifespan of mitochondrial dysfunction postmitotic cells.

Curcumin exhibits a biphasic dose-response pattern regarding the PoMiCL of wildtype yeast strain, with the most potent anti-aging effects observed at a low concentration (25 μM), while its anti-aging activity diminishes at higher concentrations (50 μM, 100 μM, and 200 μM) (Figures 2D, 1C, and 1D). Interestingly, there was no discernible hormetic effect of curcumin on the PoMiCL of mitochondrial dysfunction *pet100Δ* mutant (Figures 2B and 2C). These findings suggest that the hormetic impact of curcumin on PoMiCL is contingent upon mitochondrial functionality. To further validate this, we investigated the effect of curcumin on another mutant cells with mitochondrial dysfunction. *RMD9* (*YGL107C*) plays a pivotal role in respiration and mitochondrial genome stability ^46,47^, controlling the expression of critical mitochondrial genes, including COX2, CYTB, and ATP6, which are integral components of the OXPHOS system. Conducting PoMiCL experiments on *rmd9Δ*, along with *pet100Δ* and wildtype strains, we observed a similar influence of curcumin on the PoMiCL of *rmd9Δ* mutant, similar to that observed in *pet100Δ* mutant (Figures 2B and 2C). Our findings indicate that in postmitotic cells afflicted with mitochondrial dysfunction, curcumin (i) reverse cellular aging and prolongs lifespan, (ii) exhibits enhanced anti-aging activity with increasing concentrations, and (iii) lacks a hormetic effect on cellular lifespan.

By carefully scrutinizing all the data, we noticed that curcumin’s effective anti-aging concentrations (50 μM, 100 μM, and 200 μM) in mitochondrial mutants (Figures 2B and 2C) correspond to hormetic concentrations in wildtype cells (Figure 2D). Conversely, the curcumin concentration (25 μM) effective at promoting anti-aging in wildtype cells (Figure 2D) fails to enhance the survival of mitochondrial mutants (Figures 2B and 2C). This analysis indicates that curcumin’s anti-aging properties are influenced by both mitochondrial function and independent mechanisms. As a result, in the absence of anti-aging activity dependent on mitochondria, mitochondrial mutants experience a decreased capacity to extend cellular lifespan for an extended duration, in contrast to wildtype postmitotic cells, where both mitochondrial-dependent and independent mechanisms come into play.

### Curcumin improves mitochondrial function, leading to increased ATP levels and heightened oxidative stress

We further investigated mitochondrial function to unveil the anti-aging properties of curcumin through mitochondrial-dependent mechanisms. We assessed the impact of curcumin on mitochondrial function by measuring cellular energy levels. Wildtype cells were exposed to varying concentrations of curcumin, and we measured total cellular ATP. Our findings demonstrate that curcumin significantly increases ATP levels in cells (Figure 3A). To confirm that the rise in cellular ATP results from enhanced mitochondrial function, we examined mitochondrial-specific oxidative stress. Mitochondrial ATP synthesis is closely linked to the generation of reactive oxygen species (ROS) in cells ^48–51^. Superoxide dismutase 2 (*SOD2*) is a highly conserved mitochondrial antioxidant enzyme that protects against oxidative damage induced by ROS ^48–51^. In our yeast strain background, we initially tested whether *SOD2* is involved in oxidative stress. We exposed wildtype and *sod2Δ* deletion cells to the oxidative stress-inducing chemical hydrogen peroxide (H_2_O_2_) and assessed their growth. Our results indicated that the *sod2Δ* mutant was more sensitive to oxidative stress compared to the wildtype (Figure 3B). It is known that oxidative stress is associated with an increase in cellular aging ^48–51^. We also observed that the PoMiCL of *sod2Δ* mutant was shorter than that of wildtype (Figure S2). These findings established that *sod2Δ* mutant cells are more vulnerable to oxidative stress in our experimental yeast genetic background.

**Figure 3.**
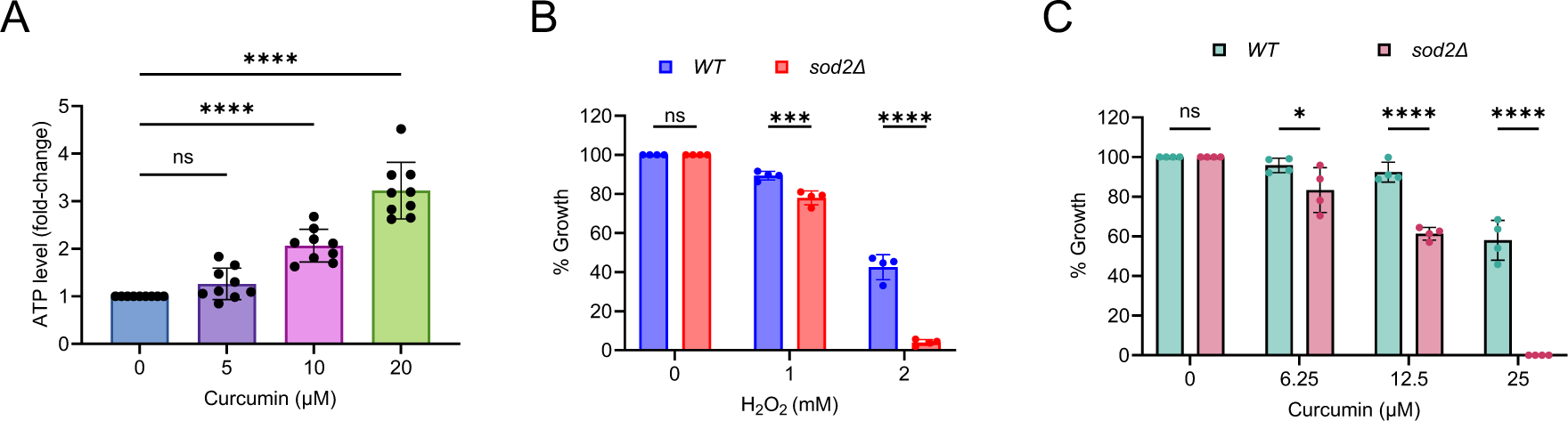
Curcumin enhances ATP levels and oxidative stress. (A) ATP analysis of yeast wildtype cells treated with indicated concentrations of curcumin for one hour. The data are presented as means ± SD (n=9). ****P < 0.0001, as determined by an ordinary one-way ANOVA followed by Dunnett’s multiple comparisons test. (B and C) Growth assay of yeast wildtype cells exposed to (B) hydrogen peroxide (H_2_O_2_) and (C) curcumin at the indicated concentrations for 72 Hours. Growth was normalized to the untreated control. The data are presented as means ± SD (n=4). Statistical significance was determined as follows: *P < 0.05, ***P < 0.001, ****P < 0.0001, and ns (non-significant) based on a two-way ANOVA followed by Šídák’s multiple comparisons test.

To further investigate whether curcumin induces oxidative stress, we examined the growth of *sod2Δ* mutant cells compared to wildtype cells when exposed to curcumin. Our results showed that the growth of curcumin-treated *sod2Δ* mutant was lower than that of wildtype cells (Figure 3C). This outcome suggests that curcumin indeed induces oxidative stress, making *sod2Δ* cells more sensitive to its effects. These results align with previous findings that have demonstrated increased ROS production following curcumin treatment in cells ^40^. Therefore, our results indicate that curcumin enhances ATP levels but also induces oxidative stress in cells. While the increase in energy may be associated with an extended cellular lifespan, the associated oxidative stress may counteract the beneficial effects of curcumin, eliciting a hormetic response in the cells.

### Curcumin inhibits TORC1, preventing accelerated aging in postmitotic cells with mitochondrial dysfunction

Target of Rapamycin Complex 1 (TORC1) is a highly conserved master regulator of cellular metabolism ^52–55^. It plays a positive role in anabolic processes while negatively impacting catabolic processes. TORC1 is a critical target for extending cellular lifespan, with its inhibition having been shown to increase healthspan and lifespan in a range of organisms, from yeast to animals ^20,33^. TORC1 senses nutrients and is closely linked to the growth and proliferation of cells. Previously, we discovered that inhibiting TORC1 can prevent the accelerated cellular aging associated with mitochondrial dysfunction ^44^. Interestingly, mutants with mitochondrial dysfunction were found to be resistant to rapamycin-induced cellular growth inhibition ^44^. Furthermore, mutants such as *pet100Δ* and *rmd9Δ* also showed resistance to the growth-inhibiting effects of curcumin (Figure 4A). These results suggest that curcumin inhibits TORC1 activity, thereby preventing accelerated aging in cells with mitochondrial function deficiencies. To confirm this, we examined the effect of curcumin on TORC1 activity by monitoring the phosphorylation of the Sch9 substrate through western blot analysis ^56^. Wildtype cells were treated with varying concentrations of curcumin, and Sch9 phosphorylation was assessed. We observed that curcumin efficiently inhibited TORC1 activity, similar to rapamycin treatment (Figures 4B; S3). This result confirmed that curcumin indeed inhibits TORC1 activity.

**Figure 4.**
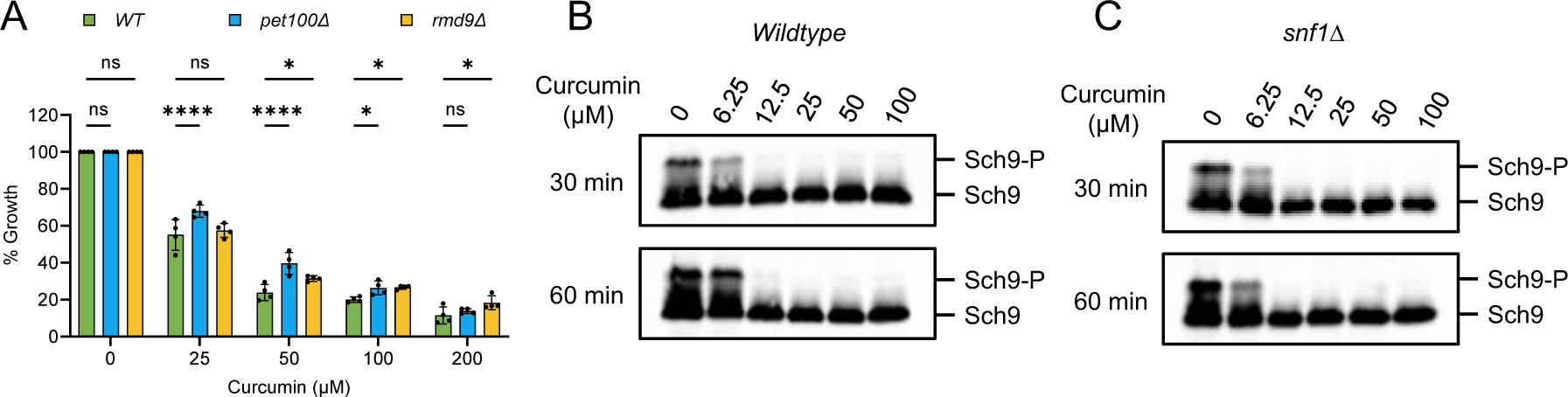
Curcumin inhibits the TORC1 signaling pathway. (A) The growth assay was conducted using yeast wildtype cells and mitochondrial dysfunction mutants (*pet100Δ* and *rmd9Δ*). They were treated with varying concentrations of curcumin for 72 hours. Growth rates were normalized to the untreated control. The data is presented as means ± SD (n=4). Statistical significance was determined as follows: *P < 0.05, ****P < 0.0001, and ns (non-significant) based on a two-way ANOVA followed by Dunnett’s multiple comparisons test. (B and C) Yeast exponential cultures of Sch9-6xHA-Tag were used for TORC1 activity analysis. In (B), wildtype cells were treated with different concentrations of curcumin for the indicated times. In (C), *snf1Δ* cells were similarly treated. Aliquots of the cultures were collected to prepare protein extracts, and TORC1 activity was assessed by monitoring the phosphorylation of the substrate, Sch9, through western blotting.

AMP-activated protein kinase (AMPK) is known to negatively regulate the TORC1 activity pathway ^57–59^. Notably, AMPK promotes catabolic processes while inhibiting anabolic pathways, contrasting with the TORC1 signaling pathway. In the yeast *Saccharomyces cerevisiae*, SNF1 protein kinase serves as the orthologue of the human AMPK complex ^60–62^. We were curious whether curcumin’s inhibition of TORC1 was dependent on the AMPK pathway. To investigate this, we assessed the effect of curcumin on TORC1 activity in *snf1Δ* deletion cells. We found that the inhibition of Sch9 phosphorylation was comparable in the *snf1Δ* mutant compared to the wildtype (Figure 4C). This result indicates that curcumin’s inhibition of TORC1 activity is independent of the AMPK pathway. Altogether, our findings reveal that curcumin delays cellular aging and increases lifespan by enhancing mitochondrial functions and reducing TORC1 activity. The inhibition of TORC1 activity, through a mitochondrial-independent mechanism, is crucial for extending the lifespan of postmitotic cells associated with mitochondrial dysfunction.

## DISCUSSION

Aging is an inevitable biological phenomenon that has been the focus of extensive research due to its profound impact on health and the growing global aging population ^1–4^. Understanding the mechanisms underlying aging and exploring strategies to mitigate its effects are essential objectives in health research. This study investigates the potential of curcumin, a polyphenolic compound derived from turmeric ^22–25^, in extending the lifespan of postmitotic cells during chronological aging, especially in the context of mitochondrial dysfunction.

### Curcumin extends postmitotic cellular lifespan

Our results demonstrate that curcumin has a significant impact on extending the lifespan of postmitotic cells during chronological aging. These findings align with previous studies that have highlighted the beneficial effects of curcumin on healthspan and cellular longevity ^26,27,37–39^. Notably, curcumin’s anti-aging effects exhibit a biphasic dose-response pattern, with lower concentrations showing greater efficacy. This hormetic effect of curcumin on cellular lifespan, observed both in yeast and human cells, adds an intriguing dimension to its potential as an anti-aging intervention.

### Mitochondrial dysfunction and postmitotic cellular lifespan

Mitochondrial dysfunction is a key factor associated with age-related diseases, including cardiovascular diseases, neurodegenerative disorders, sarcopenia, and age-related macular degeneration ^5–8^. In our study, we reveal that curcumin not only extends the postmitotic cellular lifespan of wildtype during chronological aging but also rescues the shortened lifespan of yeast cells with mitochondrial dysfunction. This is a critical finding as it suggests that curcumin may have therapeutic potential in mitigating the effects of mitochondrial dysfunction, a common hallmark of aging ^63,64^.

### Curcumin’s effects on mitochondrial function

Our investigation into the mechanisms underlying curcumin’s anti-aging properties revealed that curcumin enhances mitochondrial function by increasing ATP levels. This improvement in energy production is associated with an extension of cellular lifespan. However, it’s important to note that curcumin also induces oxidative stress, potentially as a byproduct of increased ATP synthesis. The interplay between enhanced energy production and oxidative stress may contribute to the hormetic response observed with curcumin treatment.

### Inhibition of TORC1 and cellular aging

Another key mechanism identified in this study is the inhibition of TORC1 by curcumin. TORC1 is a central regulator of cellular metabolism and aging ^52–55^, and its inhibition has been linked to increased lifespan across various organisms ^20,33^. Curcumin’s ability to inhibit TORC1 activity is crucial for its anti-aging effects, especially in cells with mitochondrial dysfunction. This finding suggests that curcumin acts through multiple pathways to promote healthy aging.

### Implications for healthy aging and longevity

The results of this study have several important implications for the field of aging research. First, curcumin derived from natural sources such as turmeric-rich diets may have the potential to promote healthy aging and extend lifespan. Second, curcumin’s ability to rescue postmitotic cells with mitochondrial dysfunction highlights its relevance for age-related diseases associated with impaired mitochondrial function. Third, the hormetic response observed with curcumin treatment underscores the importance of dosage and further emphasizes the need for careful consideration when designing interventions.

In summary, our study provides valuable insights into the intricate relationship between curcumin, postmitotic cellular aging, and mitochondrial function. It demonstrates that curcumin has the potential to extend the lifespan of postmitotic cells, with its effectiveness influenced by concentration and the presence of mitochondrial dysfunction. These findings contribute to our understanding of curcumin’s role as a potential modulator of cellular aging and emphasize the importance of considering dosage and cellular context when exploring its therapeutic applications. While this research represents a significant step forward, further investigations are needed to elucidate the precise molecular mechanisms underlying curcumin’s effects on cellular lifespan and to translate these findings into potential therapeutic interventions for human aging and age-related diseases.

## METHODS

### Yeast strains and culture conditions

We utilized the prototrophic CEN.PK113-7D strain ^65^ of *Saccharomyces cerevisiae* for our experiments. Gene deletions and protein tagging were accomplished through the application of conventional PCR-based techniques ^66^. To initiate yeast experiments, strains were resuscitated from frozen glycerol stocks and incubated on YPD agar medium, which comprised 1% Bacto yeast extract, 2% Bacto peptone, 2% glucose, and 2.5% Bacto agar. Incubation was conducted at a temperature of 30°C for a duration of 2-3 days.

### Culturing human cell lines and maintenance

Our study also involved the use of the human embryonic kidney cell line HEK293, obtained from the American Type Culture Collection (ATCC) in Manassas, Virginia, United States. These cells were maintained in D10 medium, which consists of high-glucose DMEM (HyClone #SH30022.01) supplemented with 10% fetal bovine serum (FBS) from Gibco™ (#10270106) and 1% Penicillin-Streptomycin Solution from Gibco™ (#15140122). All cell cultures were nurtured in a humidified incubator under 5% CO2 at a temperature of 37°C.

### Application of chemical compounds to cell cultures

For the introduction of chemical treatments, we prepared stock solutions of curcumin and rapamycin employing dimethyl sulfoxide (DMSO) as the solvent. The final concentration of DMSO in yeast experiments was consistently maintained below 1%, while in experiments involving human cell lines, the DMSO concentration did not exceed 0.01%. This ensured that the solvent itself did not interfere significantly with our experimental outcomes.

### Postmitotic cellular lifespan analysis of yeast during chronological aging

Lifespan of yeast postmitotic cells was assessed by determining the survival in chronological aging, as previously described ^33^. Yeast cultures were grown in synthetic defined (SD) medium (6.7 g/L yeast nitrogen base with ammonium sulfate, without amino acids, and with 2% glucose) overnight at 30°C with 220 rpm shaking. The cultures were then diluted to an initial optical density at 600 nm (OD600) of approximately 0.2 in fresh SD medium to initiate the postmitotic cellular lifespan (PoMiCL) experiment. In short, cell cultures were cultivated in 96-well plates and allowed to enter the postmitotic phase at 30°C. At various chronological time points, the survival of aging postmitotic cells was determined by measuring the outgrowth (OD600nm) in YPD medium incubated for 24 hours at 30°C, using a microplate reader.

### Postmitotic cellular lifespan analysis of human cells during chronological aging

Lifespan of human postmitotic cells was assessed by determining the survival in chronological aging using propidium iodide (PI)-fluorescent-based measurement of cell death method as previously described ^41^. Briefly, 80,000 cells with PI (5 µg/ml) in a D10 were seeded into a 96-well culture plate under non-dividing conditions with varying concentrations of curcumin and incubated at 37°C with 5% CO_2_. At different chronological aging time points, PI florescence reading was taken with a microplate reader at 535 nm excitation and 617 nm emission.

### Measurement of ATP level

For the analysis of ATP (adenosine triphosphate), yeast cells were initially treated with a 5% trichloroacetic acid (TCA) solution and chilled on ice for at least 5 minutes to preserve cellular contents. Subsequently, the cells underwent washing to remove extracellular residual before being suspended in a 10% TCA solution to aid in cellular disruption for ATP extraction. Mechanical disruption, facilitated by glass beads in a bead beater, was employed to release ATP into the solution. ATP quantification was performed using the PhosphoWorks™ Luminometric ATP Assay Kit (AAT Bioquest), likely based on a chemical reaction generating light proportional to ATP concentration. To ensure accurate comparisons, ATP levels were normalized against the protein content, determined using the Bio-Rad protein assay kit.

### Oxidative stress growth sensitivity assay

The assay was conducted to investigate the impact of oxidative stress-inducing agents on cell growth using 96-well plates. Yeast cells, with an initial OD600 of approximately 0.2 in SD medium, were transferred into the 96-well plate containing serially double-diluted concentrations of hydrogen peroxide (H_2_O_2_) or curcumin. The cells were then incubated at 30°C, and their growth was monitored by measuring OD600 using a microplate reader.

### Analysis of TORC1 activity

TORC1 activity experiments were conducted as previously described ^56^. Protein samples were separated using SDS-PAGE and transferred onto nitrocellulose membranes for Western blotting. The membranes were blocked with 5% milk in TBS/0.1% Tween 20 solution. Phosphorylation of Sch9 was assessed using an anti-HA 3F10 antibody (dilution: 1:2000; Roche Life Science, USA), followed by incubation with a goat anti-rat HRP-conjugated antibody (dilution: 1:5000; Santa Cruz Biotechnology). Blots were developed using ECL Prime Western blotting detection reagent (Amersham Pharmacia Biotech, USA) and quantified using ImageJ with the iBright CL1500 Imaging System (Thermo-Scientific).

### Data quantification and statistical interpretation

Statistical analyses of the results, including calculations of mean values, standard deviations, significance testing, and graphing, were conducted using GraphPad Prism v.10 software. Data comparisons were statistically evaluated using Student’s t-tests, Ordinary One-way ANOVA, and Two-way ANOVA, followed by post hoc multiple comparison tests. In all graph plots, significance levels are represented as *P < 0.05, **P < 0.01, ***P < 0.001, and ****P < 0.0001. Data that did not reach significance is denoted as ‘ns’ for non-significant.

## AUTHOR CONTRIBUTIONS STATEMENT

AN, YZ, NAF and SY performed the experiments. MA designed, supervised the study and wrote the manuscript.

## ACKNOWLEDGMENTS

This work was supported by A*STAR Career Development Fund (C210112008), the Global Healthy Longevity Catalyst Awards grant (MOH-000758–00), Bioinformatics Institute and Genome Institute of Singapore.

## SUPPLEMENTARY FIGURE LEGENDS

**Figure S1.**
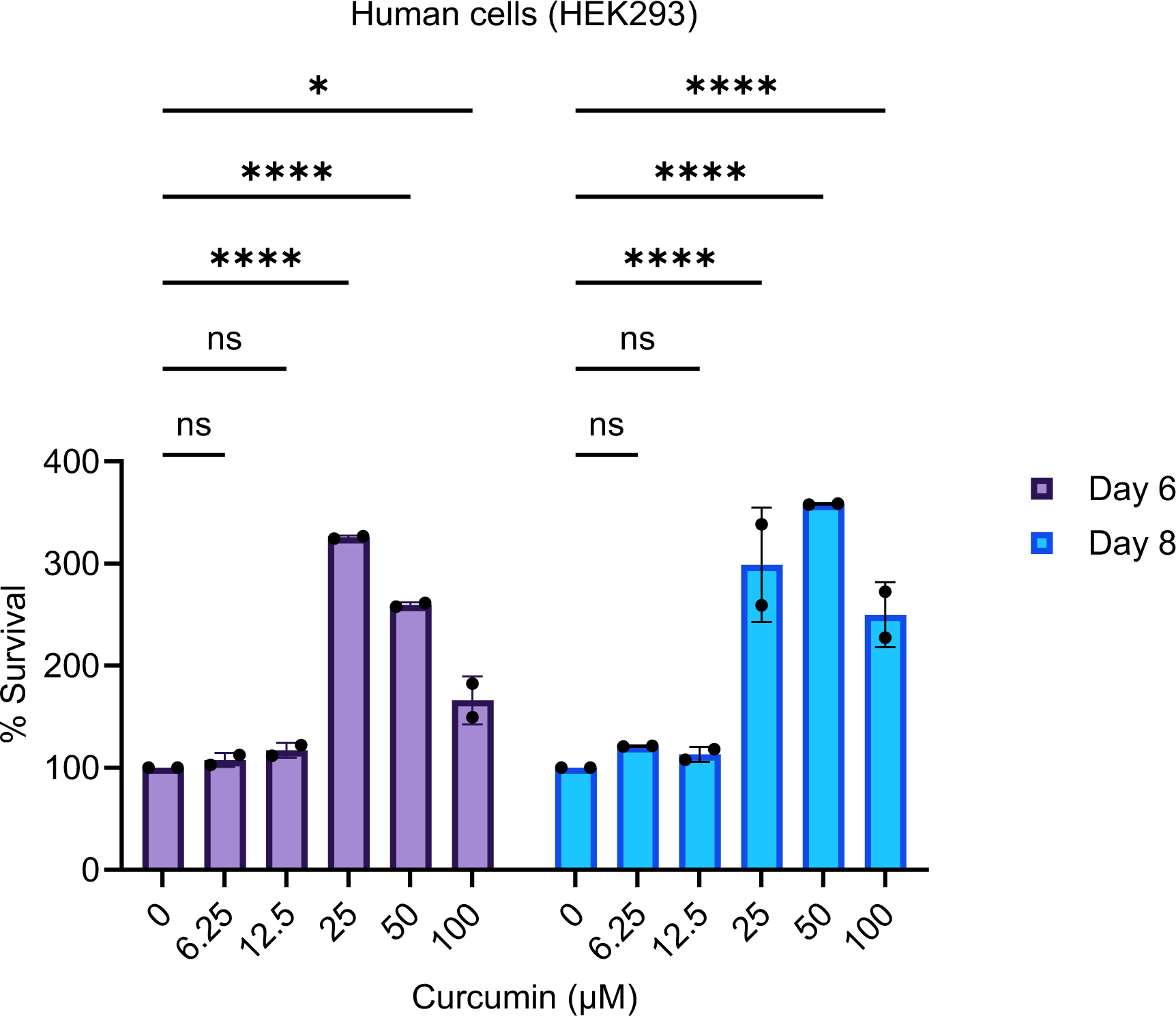
Curcumin increases the postmitotic cellular lifespan (PoMiCL) of human cells during chronological aging. The effect of curcumin treatment on the lifespan of human kidney cells (HEK293) postmitotic cells was assessed in D10 medium in a 96-well plate. The survival of aging postmitotic cells was measured using propidium iodide (PI)-fluorescent-based method for the indicated day of analysis. Data are presented as means ± SD (n=2). Statistical significance was determined as follows: *P < 0.05, ****P < 0.0001 and ns (non-significant), based on a two-way ANOVA followed by Šídák’s multiple comparisons test.

**Figure S2.**
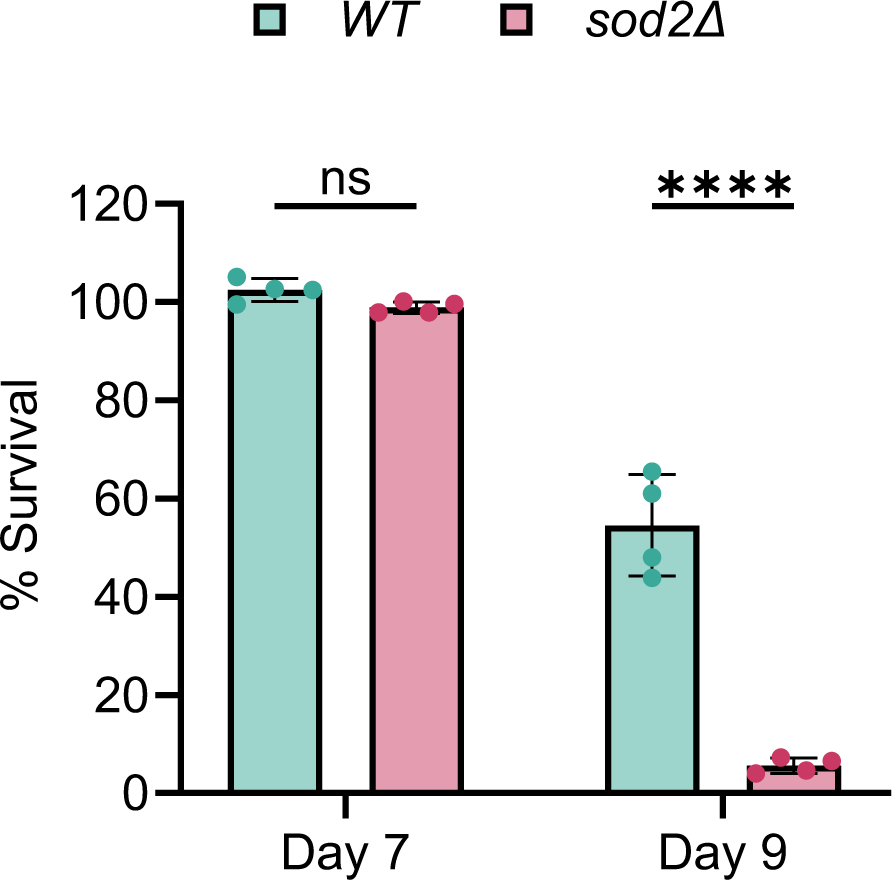
Deletion of Superoxide dismutase 2 (*SOD2*) gene decreases the postmitotic cellular lifespan (PoMiCL) lifespan of wildtype cells during chronological aging. Lifespan of yeast wildtype and *sod2Δ* deletion postmitotic cells was evaluated in an SD medium using a 96-well plate. The survival of aging postmitotic cells was measured on day 7 and day 9, relative to the outgrowth of day 3. The data are presented as means ± SD (n=4). Statistical significance was determined as follows: ****P < 0.0001 and ns (non-significant), based on a two-way ANOVA followed by Šídák’s multiple comparisons test.

**Figure S3.**
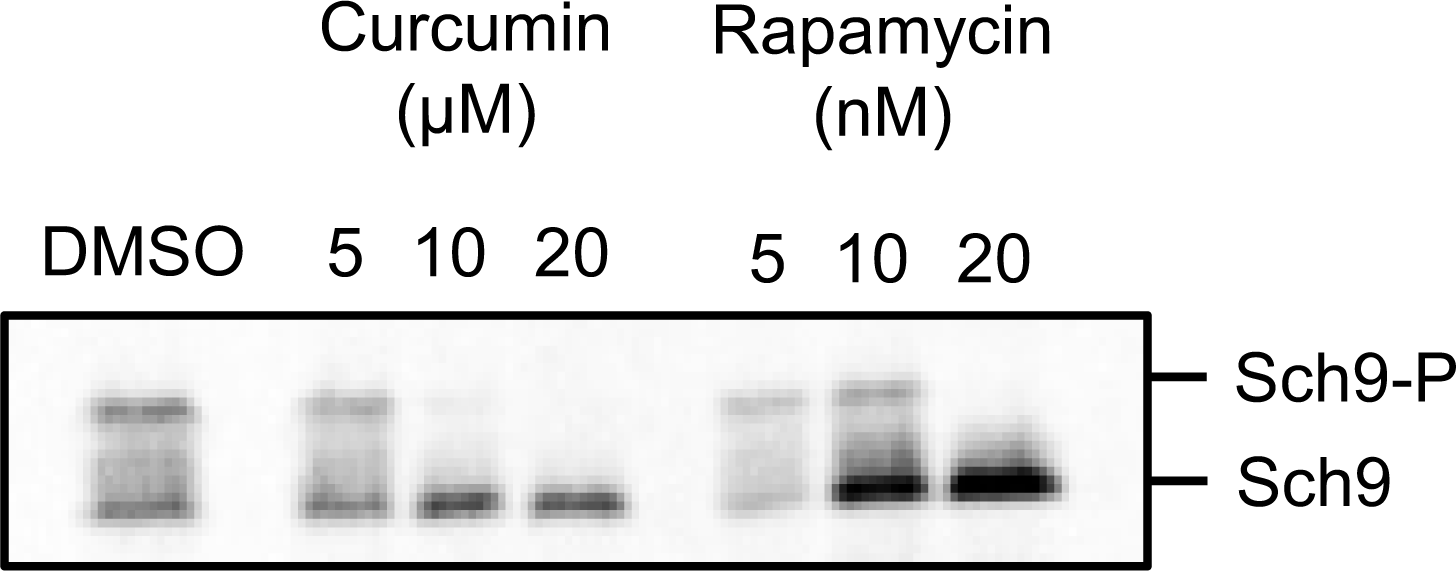
Effect of curcumin and rapamycin on TORC1 activity. Yeast exponential cultures of wildtype Sch9-6xHA-Tag cells were exposed to varying concentrations of curcumin and rapamycin for one hour. Subsequently, aliquots from these cultures were collected to extract proteins. TORC1 activity was assessed by monitoring the phosphorylation of the Sch9 substrate through western blotting.

